# Improved Mutation Detection in Duplex Sequencing Data with Sample-Specific Error Profiles

**DOI:** 10.1101/2025.07.13.664565

**Authors:** Yuhe Cheng, Shuvro P. Nandi, Luka Culibrk, Audrey Kristin, Isabella Stuewe, Shams Al-Azzam, Mia Petljak, Ludmil B. Alexandrov

**Affiliations:** Department of Cellular and Molecular Medicine, UC San Diego, La Jolla, CA, 92093, USA; Department of Bioengineering, UC San Diego, La Jolla, CA, 92093, USA; Moores Cancer Center, UC San Diego, La Jolla, CA, 92037, USA; Department of Pathology, Perlmutter Cancer Center, New York University Grossman School of Medicine, New York, NY, 10016, USA; Biomedical Sciences Graduate Program, University of California San Diego, La Jolla, CA, USA; Sanford Stem Cell Institute, UC San Diego, La Jolla, CA, USA

## Abstract

Duplex sequencing enables highly accurate detection of rare somatic mutations, but existing variant callers often rely on protocol-specific heuristics that limit sensitivity, reproducibility, and cross-study comparability. We present DupCaller, a probabilistic variant caller that builds sample-specific error profiles and applies a strand-aware statistical model for mutation detection. Across 50 synthetic datasets, DupCaller identified 1.25-fold more single-base substitutions (SBSs) and 1.41-fold more indels than a state-of-the-art method, while exhibiting equal or better precision. In three duplex-sequenced cell lines treated with aristolochic acid, it recovered expected mutational signatures while detecting 3.5-fold more SBSs and 2.8-fold more indels. In 93 tissue samples— including neurons, cord blood, sperm, saliva, and blood—DupCaller showed consistent gains, detecting 1.21- to 2.7-fold more mutations. Sensitivity scaled with sample duplication rate, yielding approximately 1.5-fold more mutations under optimal conditions and over 3-fold more in low-duplication samples where other tools falter. These results establish DupCaller as a robust and scalable solution for somatic mutation profiling in duplex sequencing across diverse biological and technical contexts.

## INTRODUCTION

Spontaneous and environmentally induced mutagenesis have gained attention for their role in the development of various human diseases^1-3^. To detect somatic mutations, studies commonly use traditional short-read sequencing of DNA from clonally expanded cells or *in vitro* amplified DNA, where multiple sequencing rounds help distinguish true mutations from random errors. Although traditional short-read sequencing introduces random errors every 1,000 to 10,000 sequenced base-pairs^4^, repeated sequencing of the same genomic region enables error correction. Because clonal expansion or amplification generates multiple copies of the same DNA molecule, true somatic mutations appear consistently across sequencing reads, whereas errors occur only sporadically. This approach has been used by a myriad of studies to sequence cells that have endogenously expanded within an individual into macroscopic normal clones^5-8^, precancerous lesions^9^, and invasive cancers^10-12^. Additionally, traditional sequencing has also been applied to *in vitro* expanded single cell derived clones or on whole-genome amplified DNA from individual cells^13,14^. Importantly, in all cases, the identified somatic mutations predominately reflect those present in the genome of a single cell that was subsequently amplified, either naturally or through experimental manipulation.

Investigating mutagenesis in healthy tissues necessitates profiling mutations at ultra-low allelic fractions, as each cell harbors a distinct set of somatic mutations^2,15^. Unfortunately, the high error rate of traditional short-read sequencing renders it inadequate for this task^16^. While single-cell DNA sequencing^17-20^ or single-cell clonal expansion^13,14,21^ are possible, scaling these methods is labor-intensive, costly, and requires sequencing many cells to achieve a representative view of overall tissue mutagenesis.

Error-corrected duplex sequencing has emerged as a powerful alternative for profiling somatic mutations and mutation rates in healthy tissues.^22^ By leveraging the inherent complementarity of genomic DNA and randomly barcoded adapters during library preparation^23,24^, duplex sequencing greatly enhances base calling accuracy, enabling the detection of somatic mutations at ultra-low allelic fractions^23,24^. In this method, two random barcodes are attached to both ends of the DNA template (**Figure 1*a***), ensuring all reads from the same fragment share the same barcode. This approach enables reliable discrimination between true mutations and sequencing errors: genuine mutations are consistently supported by reads from both DNA strands, whereas errors introduced during sequencing or amplification generally appear on only one strand or in a small subset of reads. Computational analysis of duplex sequencing data typically begins by generating single-strand consensus sequences (SSCS) by grouping reads originating from the same DNA strand. These are then paired and merged into duplex consensus sequences (DCS), which are used for accurate variant calling (**Figure 1*a***). Following the emergence of duplex sequencing, numerous short-read sequencing protocols have been developed to lower error-rates and reduce sample costs. NanoSeq^15^ is currently the state-of-the-art method for detecting somatic mutations, achieving an exceptionally low error rate of just five errors per billion base pairs sequenced (*i.e.,* 5 x 10^-9^ errors per base-pair).

**Figure 1.**
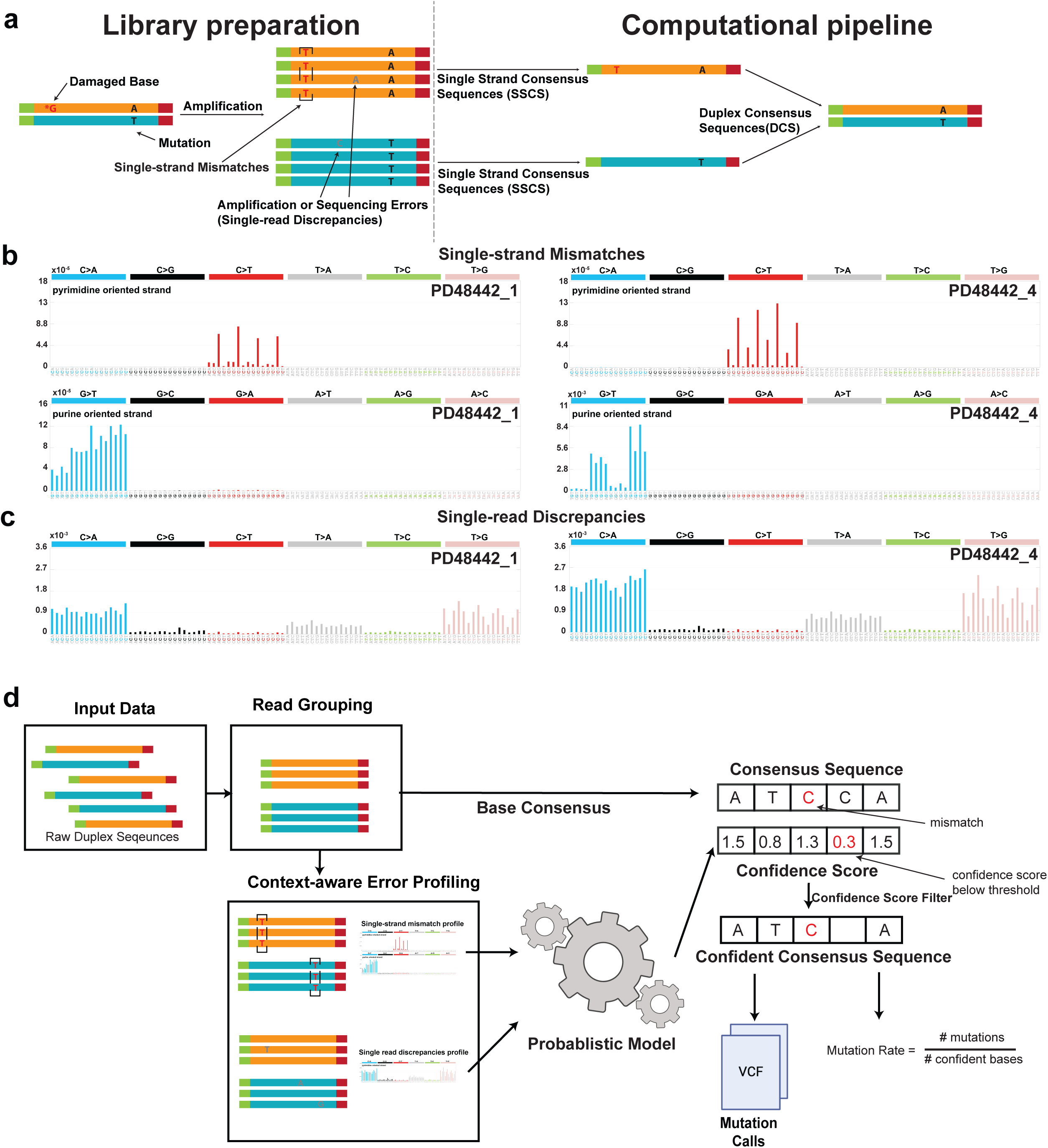
Illustration of duplex sequencing, sample-specific error profiles, and DupCaller algorithm. ***(a)*** Simplified illustration of the duplex sequencing library preparation process *(left)* and a representative computational pipeline for mutation calling *(right)*. DNA damage, such as oxidized guanine, can lead to single-strand mismatches that appear as thymine after amplification (highlighted in red). Single-read discrepancies introduced during amplification or sequencing are also shown (gray letters). Computationally, reads are collapsed into single-strand consensus sequences (SSCSs), which enable the detection of single-strand mismatches. SSCSs from complementary strands are then combined to form duplex consensus sequences (DCSs) for high-confidence variant detection. ***(b–c)*** Frequencies of mismatch and discrepancy types shown in trinucleotide context for two replicates of PD48442 NanoSeq data (PD48442_1 and PD48442_4). *(b)* Single-strand mismatches and *(c)* single-read discrepancies are presented in full trinucleotide context, incorporating the mutated base along with the 5′ upstream and 3′ downstream bases. Single-strand mismatches are further oriented based on whether the mutated base is a pyrimidine or purine, reflecting strand-specific DNA damage. ***(d)*** Overview of the DupCaller algorithm. Raw duplex reads are grouped by shared molecular barcodes to generate SSCSs, which are then merged into DCSs. Context-aware error profiles—specific to each sample—are constructed for both single-strand mismatches and single-read discrepancies, stratified by trinucleotide context and base class (purine versus pyrimidine). These profiles inform a probabilistic model that assigns confidence scores to each base. Bases with low confidence scores are filtered, producing a high-confidence consensus sequence. These sequences are used for mutation calling and estimation of mutation rates, with final variants reported in VCF format.

Most duplex sequencing datasets are processed using lab-specific pipelines with custom filtering and tuned parameters, making cross-study comparisons difficult^15,25-27^. For instance, NanoSeq data are typically analyzed using the NanoSeq pipeline, which strictly adheres to the original duplex sequencing framework (**Figure 1*a***)^24^ — assuming strand symmetry and applying heuristic cutoffs that may exclude a substantial fraction of genuine somatic mutations along with noisy data. To address these limitations, we developed Duplex-based Variant Caller (DupCaller), a computational tool that standardizes mutation calling and processing of duplex sequencing data. DupCaller employs a strand-specific probabilistic model to detect somatic mutations and quantify mutational burden. Validation against NanoSeq protocol-specific software using synthetic and multiple real datasets demonstrate DupCaller’s ability to identify substantially more somatic mutations while effectively minimizing potential artifacts.

## RESULTS

### Sample-specific Error Profiles in Duplex Sequencing Data

Errors in duplex sequencing data can be broadly categorized as single-strand mismatches or single-read discrepancies. Single-read discrepancies are sporadic, low-frequency errors that typically arise from DNA polymerase mistakes during PCR amplification^28^, sequencing artifacts^29^, or other stochastic processes (grey letters in **Figure 1*a***). These discrepancies affect only one read within a strand family and are not reproducible across other reads from the same DNA molecule. As a result, they generally fail to form consensus bases in single-strand consensus sequences (SSCSs), and—if uncorrected—can contribute to false-positive variant calls. On the other hand, single-strand mismatches typically arise from DNA damage or lesions that affect only one strand prior to amplification^30^. These errors are propagated during PCR, leading to consistent changes across reads from the same strand. As a result, they produce concordant SSCSs but generate mismatches when SSCSs from opposite strands are compared, leading to discordant duplex consensus sequences (DCSs; red letters in **Figure 1*a***). If not properly accounted for, single-strand mismatches can be misinterpreted as true somatic mutations. Notably, single-strand mismatches retain the strand specificity of the DNA damage, which can be inferred from the read strand. As illustrated in **Figure 1*a***, the concordant SSCSs and discordant DCS indicate the presence of a mismatch, DNA damage, or other strand-specific aberration, even though the specific damaged base cannot be identified without prior information.

In duplex sequencing, single-read discrepancy error profiles can be built by comparing original sequencing reads to their SSCSs, revealing errors that appear only in individual reads. Similarly, single-strand mismatch error profiles can be built by comparing SSCSs from each strand to the final DCS, capturing strand-specific errors such as those caused by DNA damage. Since purine and pyrimidine bases have different susceptibilities to DNA damage^30^, single-strand mismatches can also be oriented based on the pyrimidine or purine base of the Watson-Crick pair, producing distinct error profiles. To illustrate this, we analyzed two previously generated NanoSeq replicates from the same cord blood sample^15^ PD48442_1 (sample ID 32268#46) and PD48442_4 (sample ID 33796#35), and calculated their sample-specific error profiles (**Figure 1*b-c***). In particular, single-strand discrepancies for each SBS-96 mutation type—which classifies single base substitutions (SBS) into 96 categories based on the pyrimidine base of the mutated base-pair and its immediate trinucleotide context^31^—revealed a profile with high rates of C>A, T>A, and T>G mutations (**Figure 1*c***). Additionally, we derived the SBS-96 error profiles for single-strand mismatches oriented based on the pyrimidine and purine base, which displayed distinctly different patterns (**Figure 1*b***). Specifically, the pyrimidine orientation was predominantly characterized by C>T mutations at CpG sites, likely resulting from the deamination of methylated cytosine^32^. In contrast, the purine orientation was mainly dominated by G>T mutations, which are likely associated with oxidative damage, such as 8-oxo-guanine formation^33^. Importantly, error profiles are sample-specific and can differ markedly in both patterns and magnitudes, as demonstrated by the two replicates of sample PD48442_1 (**Figure 1*b-c***). Clear differences in the error profiles are evident for both pyrimidine- and purine-oriented single-strand mismatches. Moreover, the error profile of the purine-oriented strand shows almost a two-order-of-magnitude difference between replicates (**Figure 1*b***).

### Overview of the DupCaller Algorithm

Current duplex sequencing analysis software generally relies on a predefined minimum number of reads per strand to generate consensus sequences^15,24,25^. For instance, the NanoSeq analysis software^15^ uses a default threshold of at least two reads per strand to identify mutations, discarding read families supported by only one strand or those where a single read supports one of the strands. While this approach effectively reduces artifacts, it also excludes relevant sequencing data with a lower duplication rate, potentially leading to the loss of genuine somatic mutations during analysis. To address this limitation and integrate sample-specific error profiles into mutation calling, we developed a probabilistic model-based variant caller for duplex sequencing data, termed Duplex-based Variant Caller (DupCaller; **Figure 1*d***). For SBSs, DupCaller uses a probabilistic model that derives sample-specific SBS-96 error profiles for single-strand discrepancies as well as for single-strand mismatches in purine and pyrimidine orientation (**Methods**). For short insertions and deletions (indels), the tool calculates error rates based on the homopolymer lengths preceding and following each location (**Methods**). By accounting for error rate variations across trinucleotide contexts and homopolymer lengths, DupCaller assigns higher confidence to mutations in rarely damaged contexts, thereby increasing the number of mutations that can be detected while maintaining high accuracy.

### Benchmarking DupCaller Using Synthetic Data

Benchmarking and comparing the performance of duplex sequencing variant calling pipelines is challenging due to the difficulty in validating mutations, since most mutations originate from a single cell and cannot be directly confirmed. To address this, we developed an *in-silico* germline duplex mutation spike-in approach to create synthetic datasets with known ground truth using previously generated data from the NanoSeq protocol^15^. In particular, we used both diluted and undiluted cord blood NanoSeq data from two donors (PD47269, *donor A*, and PD48442, *donor B*). In principle, the diluted sequencing data is used to detect non-clonal somatic mutations, while the undiluted data serves as a matched-normal to filter out germline variants and clonal mutations. From the undiluted datasets, we computationally extracted read groups containing germline mutations present in *donor A* but absent in *donor B* and designated these as the ground truth (**Figure 2*a***; **Methods**). Next, we computationally integrated randomly selected read groups from the ground truth into the undiluted data of *donor B*, introducing known mutations. It should be noted that this approach has at least two limitations: *(i)* true mutations originating from *donor B* will be misclassified as false positives, reducing precision estimates; and *(ii)* the introduced mutations are germline variants, which necessitates customizing the analysis pipelines, as most pipelines rely on germline filters during mutation calling. Using NanoSeq of *donor A* (PD47269) and five replicates from *donor B* (*PD48442)*, we created 25 synthetic SBS datasets with ground truth sizes ranging from 100 to 5,000 SBSs and 25 synthetic indel datasets with ground truth sizes ranging from 10 to 500 indels, covering a range of real-world scenarios. We then compared DupCaller’s performance on these 50 datasets against the NanoSeq analysis software from the original publication^15^.

**Figure 2.**
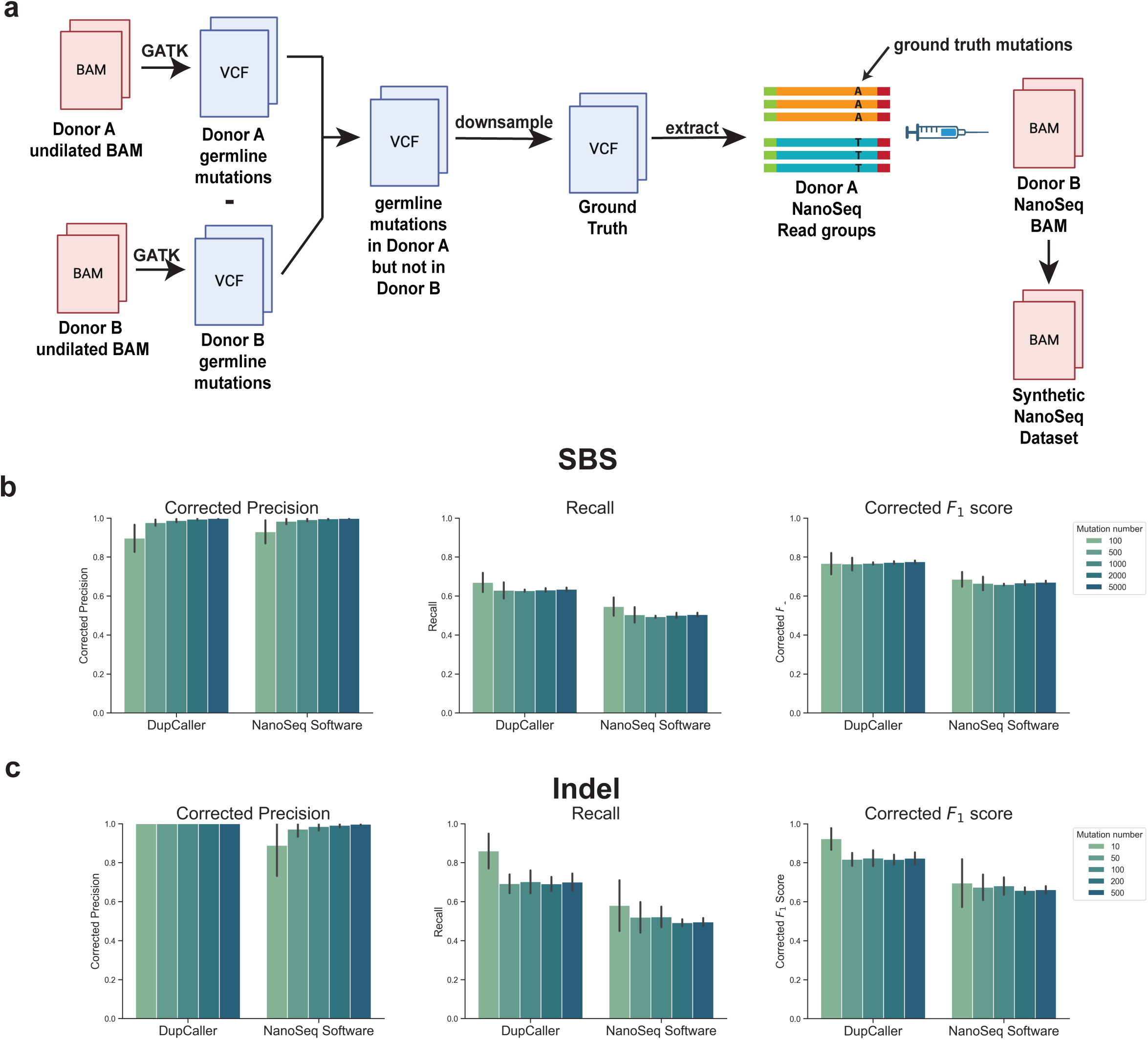
Benchmarking DupCaller using synthetic duplex sequencing datasets. ***(a)*** Schematic of the *in silico* synthetic data generation approach used to create duplex sequencing datasets with known ground truth. To enable controlled benchmarking, we developed a germline mutation spike-in strategy using previously generated NanoSeq data from two donors: PD47269 (*donor A*) and PD48442 (*donor B*). Germline variants unique to *donor A* were identified by comparing undiluted BAMs. These unique variants were downsampled and defined as ground truth. We then extracted read groups carrying these mutations from diluted *donor A* NanoSeq data and computationally spiked them into diluted *donor B* BAM file, ensuring that the loci were well covered and that no supporting reads were present in *donor B*. This resulted in a synthetic NanoSeq dataset containing known mutations at predefined locations. ***(b–c)*** Benchmarking results comparing DupCaller and the NanoSeq analysis software across 25 synthetic datasets of 5 mutation counts, 5 dataset per mutation count. Metrics include recall, corrected precision, and corrected F_1_ score for both single base substitutions (SBS; *b)* and insertions/deletions (indels; *c)*. Precision was corrected to account for potential true somatic mutations present in *donor B* that were not part of the defined ground truth. DupCaller consistently demonstrated higher recall and F_1_ scores, especially at lower mutation burdens, while maintaining comparable or improved precision. Bars are color-coded by the number of spiked-in mutations. The bar plot for each mutation count represents the average metrics per mutation count, while the error bars represent the standard deviation.

For somatic substitutions, DupCaller’s precision-recall curve outperformed the NanoSeq analysis software in 24 out of 25 synthetic datasets (**Figure S1**). DupCaller also achieved higher F_1_ scores—the harmonic mean of precision and recall, reflecting overall mutation-calling performance—in 21 datasets (**Figure S2*a***) and consistently demonstrated superior recall across all comparisons (**Figure 2*b***). In the four datasets where DupCaller showed slightly lower F_1_ scores, each contained fewer than 100 ground truth mutations. One possible explanation is DupCaller’s increased sensitivity in detecting more mutations originating in *donor B*. Since DupCaller identifies more sites than NanoSeq for the same sample, it may capture more true mutations in *donor B* that are mistakenly labeled as false positives in the benchmark. To account for this, we estimated the expected number of mutations based on the known mutation rate from cord blood colony whole-genome sequencing of *donor B*^15^ and the number of sites called by each pipeline (**Methods**). After this correction, DupCaller’s precision matched that of NanoSeq across all samples, and the corrected F_1_ scores showed that DupCaller outperforms NanoSeq in all 25 datasets (**Figure 2*b***). For somatic indels, DupCaller’s precision-recall curve exceeded NanoSeq analysis software’s precision-recall point in all 25 datasets (**Figure S3**), as well as its corrected F_1_ scores outperformed NanoSeq analysis software in all 25 datasets (**Figure 2*c*; Figure S2*b***).

Importantly, DupCaller maintained high corrected precision (>0.90) in all samples with more than 500 SBSs and 100% corrected precision in all indel datasets (**Figure 3*c***). To investigate sources of false negatives—missed genuine somatic mutations—we analyzed their distribution and contributing factors. We found that 19% of all missed SBSs occurred within the first 7 bases of the 5’ or 3’ ends—regions typically excluded by both DupCaller and NanoSeq analysis software due to elevated error rates in duplex sequencing data. In contrast, only 3% of missed indels fell within these regions, likely due to alignment artifacts near read ends. Additionally, 25% of missed SBSs and 27% of missed indels were associated with low coverage (<10 reads) in the undiluted control. The causes of the remaining false negatives remain unclear.

**Figure 3.**
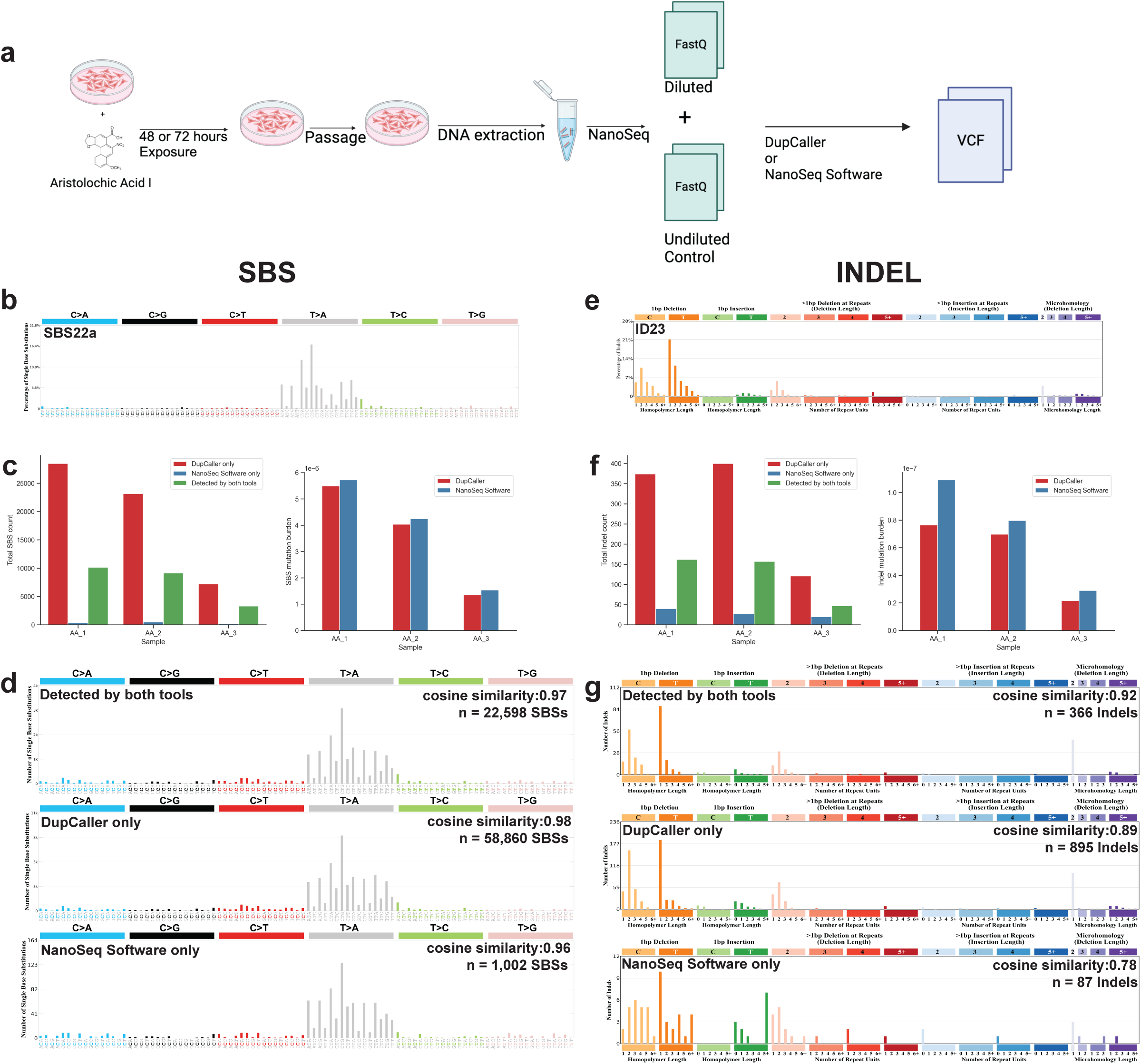
DupCaller benchmarking using aristolochic acid exposure in a cell line model. ***(a)*** Experimental workflow. HepG2 cells were exposed to aristolochic acid I (AA) for 48 or 72 hours, followed by cell passage, DNA extraction, and duplex sequencing using the NanoSeq protocol. Both diluted and undiluted libraries were generated for each replicate and analyzed using DupCaller or the NanoSeq analysis software. ***(b)*** The expected AA substitution signature SBS22a, characterized primarily by T>A substitutions, commonly observed in AA-exposed tumors. ***(c)*** Comparison of single base substitution (SBS) call sets across three AA-treated samples. Bar plots show the number of SBSs detected exclusively by DupCaller (red), exclusively by NanoSeq analysis software (blue), or by both tools (green), along with the estimated SBS burden per sample. ***(d)*** SBS-96 mutation spectra for each call set category. Cosine similarities with SBS22a are shown along with the total number of SBSs in each category across all replicates. ***(e)*** The expected AA indel signature ID23, characterized by single-base deletions of cytosine and thymine, typical of AA-induced mutagenesis. ***(f)*** Comparison of indel call sets and estimated indel burdens across the same three AA-treated samples, using the same color coding as in panel *(c)*. ***(g)*** ID-83 mutation spectra for each indel call set category. Cosine similarities with ID23 are shown along with the total number of indels in each category.

In summary, based on our synthetic benchmark, DupCaller demonstrated better sensitivity on synthetic datasets, consistently outperforming NanoSeq analysis software by detecting 25% more SBSs and 41% more indels on average.

### Benchmarking DupCaller Using Experimental Data

As previously discussed, benchmarking duplex sequencing variant-calling pipelines is challenging because most detected mutations occur in individual cells and cannot be independently validated. One approach to overcome this limitation is to expose cell lines to a mutagen with a known mutational signature and assess whether each computational pipeline detects the expected mutagenic pattern. To validate our approach, we conducted an *in vitro* experiment in which the human hepatocellular carcinoma cell line HepG2 was exposed to aristolochic acid I at three different dose-time combinations (**Figure 3*a*; Methods**). Aristolochic acid I is a potent mutagen known to induce substitution signature SBS22a, characterized primarily by T>A mutations (**Figure 3*b***)^34,35^, and indel signature ID23, characterized by one base-pair deletions of cytosine and thymines (**Figure 3*e***)^36^. For each condition, DNA was extracted and subjected to NanoSeq, with both diluted samples and undiluted controls generated. Somatic mutations were independently identified from the sequencing data using both the NanoSeq analysis software and DupCaller.

In line with results from benchmarking with synthetic data, DupCaller identified more somatic mutations than the NanoSeq analysis software—detecting 3.5 times more SBSs and 2.8 times more indels across the three conditions—while yielding comparable mutation rate estimates (**Figure 3*c***). Importantly, almost all mutations detected by the NanoSeq analysis software were also identified by DupCaller (**Figure 3*c&f***). Moreover, SBSs (*n*=22,598) and indels (*n*=366) detected by both DupCaller and NanoSeq analysis software showed a high cosine similarity of 0.97 to SBS22a (**Figure 3*d***) and 0.92 to ID23 (**Figure 3*g***), respectively. Notably, the SBSs uniquely detected by DupCaller (*n*=58,860) and those identified only by NanoSeq (*n*=1,002) both showed high cosine similarity to SBS22a (>0.96), indicating that DupCaller captures a substantial proportion of true SBSs while missing only a small subset of mutations detected by NanoSeq analysis software (**Figure 3*d***). Interestingly, indels detected exclusively by DupCaller (*n*=895) exhibited a cosine similarity of 0.89 to ID23 (**Figure 3*g***). In contrast, indels identified solely by the NanoSeq analysis software (*n*=87) displayed a markedly different pattern, with a cosine similarity of 0.78 and unexpected peaks of single-base thymine insertions or deletions at long homopolymers (**Figure 3*g***). These findings suggest that DupCaller is more effective at distinguishing true indels from potential sequencing artifacts, whereas the NanoSeq analysis software may yield a small number of potentially artifactual indels at long homopolymers. The >3-fold increase in mutation counts is particularly striking, and one likely contributor to this elevation is the lower duplication rate observed in these datasets (Figure S4), which results in smaller read family sizes compared to standard NanoSeq^15^. This underscores DupCaller’s strong performance even under suboptimal duplication conditions.

### Benchmarking DupCaller Using Tissue Data

We further applied DupCaller to NanoSeq data from 17 neuron samples and 7 cord blood samples (one from donor PD47269 and six replicates from donor PD48442) from Abascal *et al.* ^15^ to assess its performance in both moderately and lowly mutated tissues, and to compare its results with those obtained using the NanoSeq analysis software. DupCaller captured nearly 100% of the SBSs identified by the NanoSeq analysis software in both neuron and cord blood samples, while detecting an additional 50% more SBSs in neuron samples (1.5-fold increase) and approximately 100% more in cord blood samples (2-fold increase), consistent with its high sensitivity (**Figure 4*a***). Despite identifying more SBSs, the overall mutational burden estimates for neuron and cord blood samples remained largely stable (**Figure 4*b***). This is because DupCaller also expanded the number of confidently analyzable base-pairs across the genome, effectively increasing the denominator used to normalize mutation counts. Moreover, the SBS-96 pattern of DupCaller-specific mutations closely resembled those of mutations shared between DupCaller and NanoSeq analysis software in both neuron (**Figure 4*c***) and cord blood samples (**Figure S5*a***), supporting that the additional calls represent *bona fide* somatic mutations.

**Figure 4.**
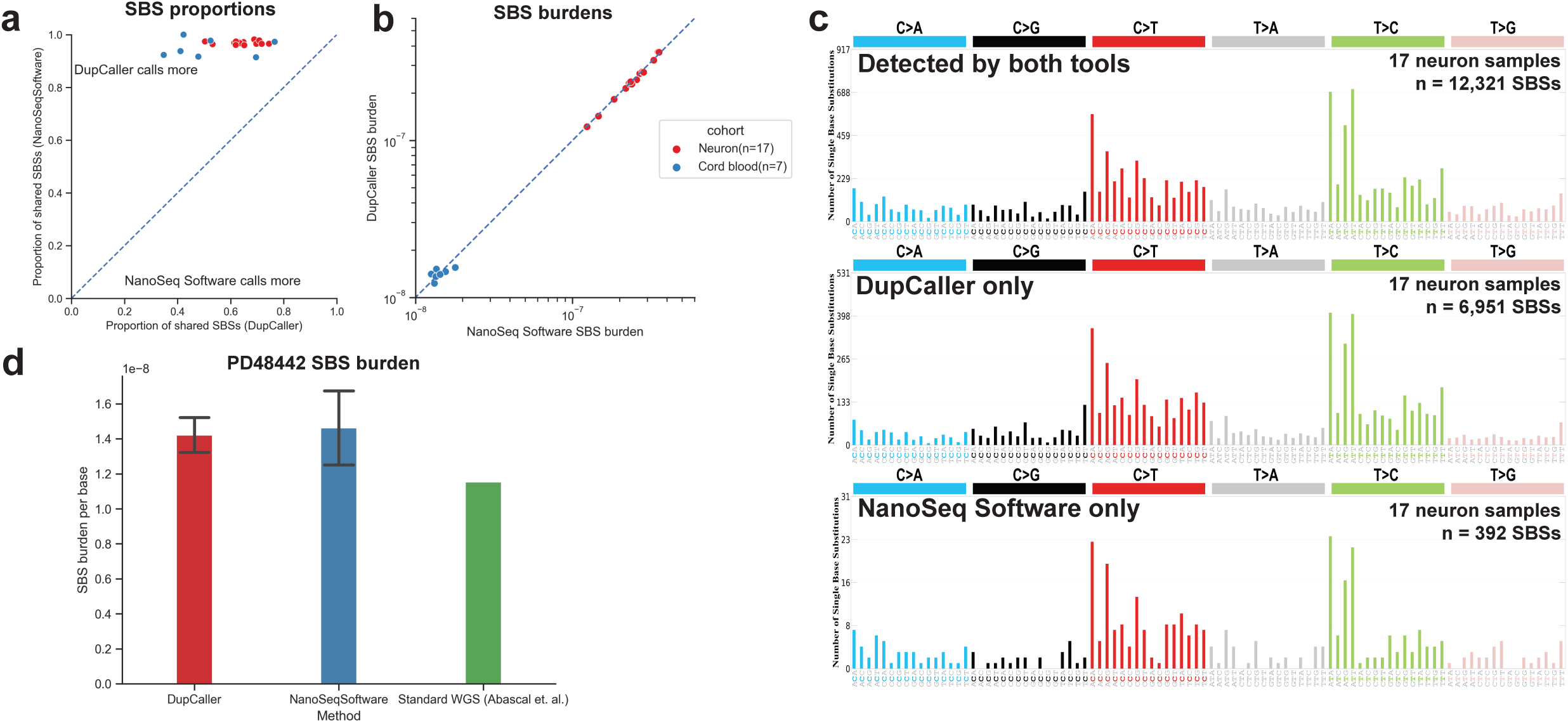
Application of DupCaller for single base substitution detection in normal tissue duplex sequencing data. ***(a)*** Proportion of shared single base substitutions (SBSs) between DupCaller and NanoSeq analysis software for each sample. The x-axis shows the proportion of shared SBSs relative to the total number of SBSs called by NanoSeq analysis software; the y-axis shows the same proportion relative to DupCaller calls. Points above the diagonal indicate more unique SBSs from DupCaller. Points below the diagonal indicate more unique SBSs from NanoSeq analysis software. **(b)** Comparison of SBS burdens per sample between DupCaller and NanoSeq analysis software. The x-axis shows SBS burden estimated by NanoSeq analysis software; the y-axis shows SBS burden estimated by DupCaller. ***(c)*** SBS-96 mutational spectra of SBSs detected by both tools (top), DupCaller only (middle), and NanoSeq analysis software only (bottom), across 17 neuron samples. The x-axis denotes the 96 trinucleotide mutation types; the y-axis represents mutation counts. ***(d)*** SBS burden estimates in PD48442 cord blood samples using DupCaller and NanoSeq analysis software, compared to standard WGS of cord blood colonies. The x-axis shows the method; the y-axis shows the estimated mutation rate of SBSs.

In neuron samples, DupCaller detected 1.4-fold more indels compared to the NanoSeq analysis software (**Figure 5*a***). In cord blood samples, DupCaller also identified 1.2-fold more indels, however, NanoSeq software unique calls were almost exclusive single-base deletions or insertions of thymine located in homopolymer regions (**Figure S5*b***) and may represent sequencing artifacts. Supporting this, DupCaller’s indel burden estimates more closely matched those obtained from whole-genome sequencing of cord blood colonies from the same donor^15^, suggesting higher accuracy in its indel detection (**Figure 5*d***). Furthermore, when examining aggregate indel patterns across all neuron samples, indels uniquely identified by the NanoSeq analysis software showed an increased frequency of single thymine insertions and deletions in long homopolymers (**Figure 5*c***).

**Figure 5.**
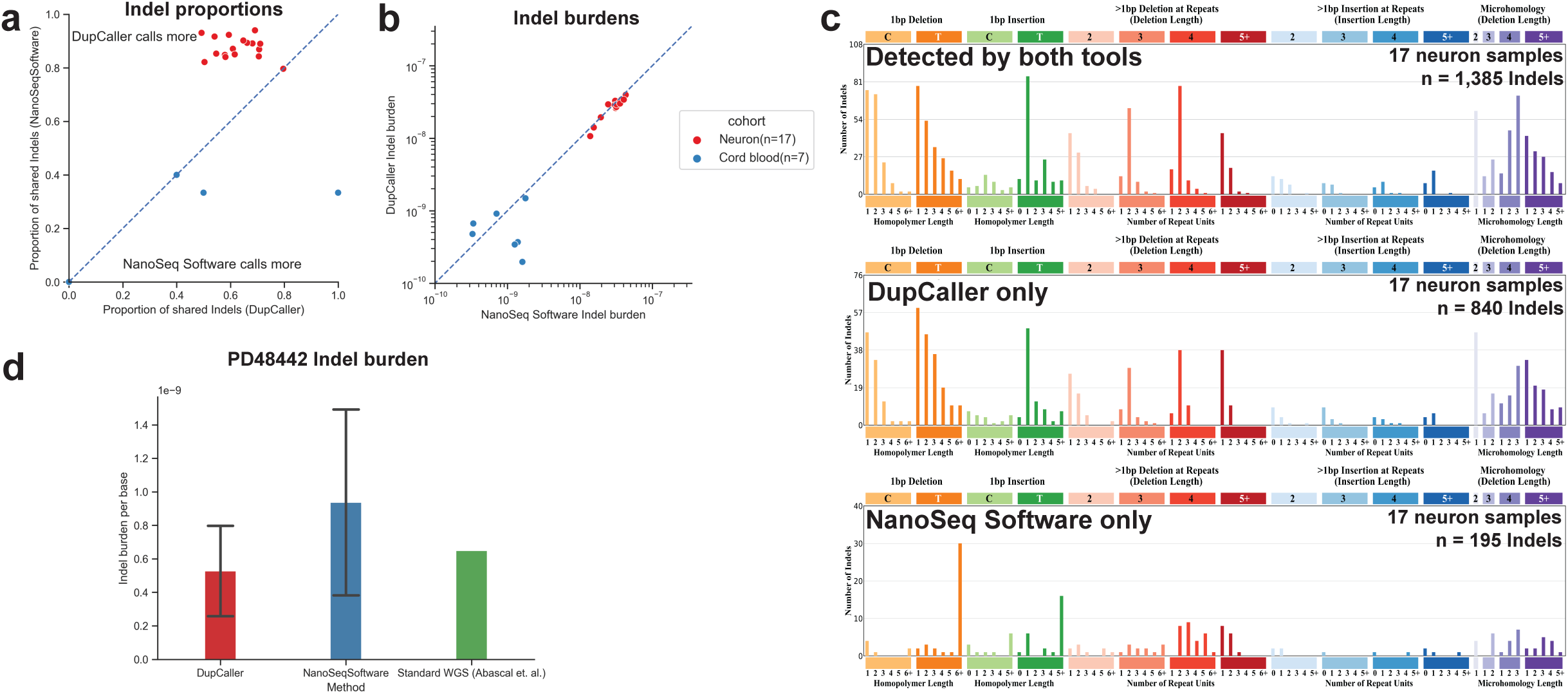
Application of DupCaller for indel detection in normal tissue duplex sequencing data. ***(a)*** Proportion of shared indel calls between the two pipelines. The format of the plot is similar to the one in Figure 4a. Samples with no shared indels are represented at the origin (x = 0, y = 0). ***(b)*** Comparison of indel burden per sample between DupCaller and NanoSeq analysis software. The x-axis shows indel burden estimated by NanoSeq analysis software; the y-axis shows indel burden estimated by DupCaller. ***(c)*** ID-83 mutational spectra of indels detected by both tools (top), DupCaller only (middle), and NanoSeq analysis software only (bottom), across 17 neuron samples. The x-axis denotes the 83 indel categories; the y-axis shows mutation counts. ***(d)*** Indel burden estimates in PD48442 cord blood samples using DupCaller and NanoSeq analysis software, compared to WGS-based estimates. The x-axis shows the method; the y-axis shows the estimated number of indels per genome.

To further validate DupCaller’s performance, we applied it to an independently published NanoSeq dataset comprising 46 sperm, 15 blood, and 8 saliva samples from 23 donors^37^. DupCaller reported SBS mutation burdens nearly identical to those generated by the original NanoSeq software (**Figure 6*a***), while identifying 1.5-fold more SBSs on average (range: 1.21–1.92; **Figure 6*b***). Notably, the SBS96 mutational spectrum of DupCaller’s call set closely resembles that of paternal *de novo* mutations^38^ (cosine similarity = 0.98; **Figure 6*c***), indicating that DupCaller maintains high precision while substantially increasing sensitivity.

**Figure 6.**
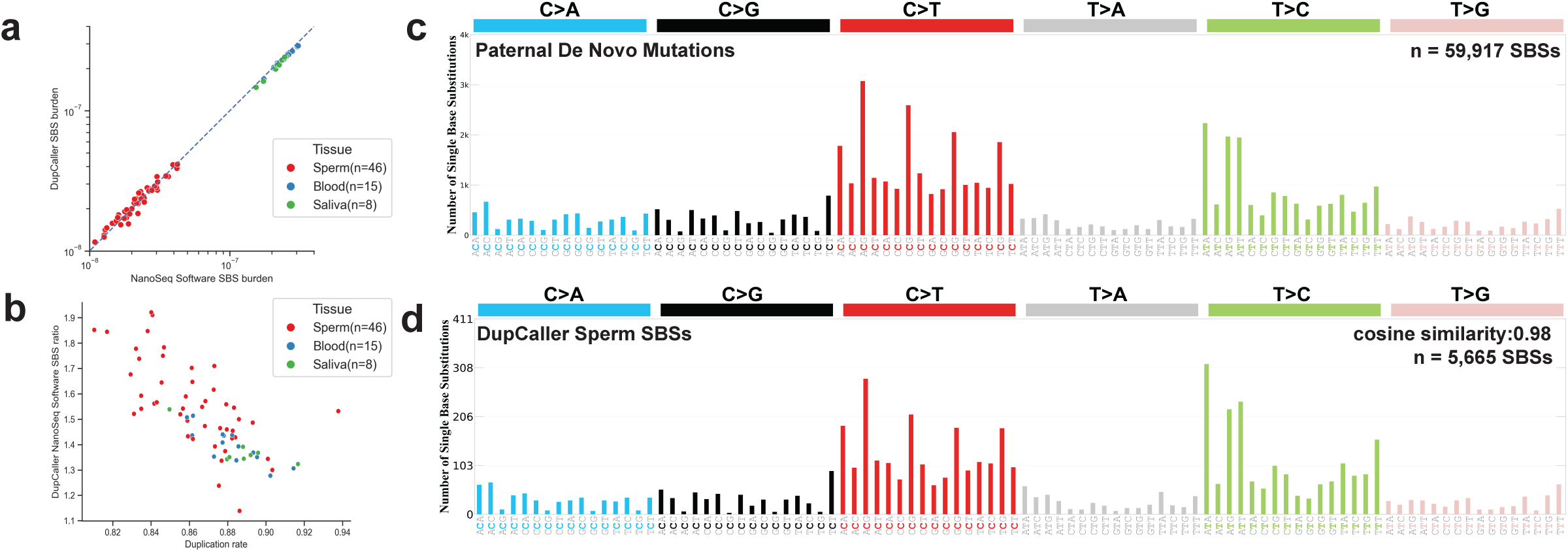
DupCaller benchmarking on NanoSeq data from sperm, blood, and saliva samples. ***(a)*** Comparison of single base substitution (SBS) burden per sample between DupCaller and NanoSeq analysis software. The x-axis shows SBS burden estimated by NanoSeq analysis software; the y-axis shows indel burden estimated by DupCaller. ***(b)*** The ratio of DupCaller and NanoSeq analysis software’s number of SBSs called versus the duplication rate of each sample. ***(c)*** SBS-96 mutational spectra of parental de novo mutations from 2,976 trios.^38^ ***(d)*** SBS calls by DupCaller in sperm samples closely resemble the paternal *de novo* mutation spectrum, with an SBS-96 cosine similarity of 0.98 based on data from 2,976 trios.

### DupCaller Gains Are Modulated by Duplication Rate

As previously shown, the increase in mutation detection by DupCaller is influenced by the duplication rate of the sample, with lower duplication rates enabling more substantial gains (**Figure 6*b***; **Figure S4**). DupCaller’s strand-specific probabilistic model allows it to recover genuine mutations from smaller read families that are typically excluded by conventional, threshold-based pipelines. For instance, in AA–treated samples with a ∼60% duplication rate, DupCaller identified over three times more mutations than the NanoSeq analysis software. In neuron samples with ∼80% duplication, the increase was more moderate at >1.5-fold, and in sperm samples with ∼90% duplication, the gain was approximately 1.35-fold. Given that the NanoSeq method estimates^15^ an optimal duplication rate of ∼81%, DupCaller is expected to yield ∼1.5-fold more mutations in well-optimized samples. Crucially, it also enables robust and accurate mutation detection in sub-optimally prepared samples, significantly expanding the utility of duplex sequencing to a broader range of experimental conditions.

### DupCaller Computational Performance

To assess computational efficiency, we benchmarked DupCaller against the NanoSeq analysis software across five key processing steps using 17 neuron samples^15^. DupCaller outperformed NanoSeq in each step, owing to its streamlined workflow and optimized algorithms (**Figure S6**). Notably, the mutation calling step—typically the most computationally demanding—was 9.8 times faster on average when run on identical hardware. Overall, DupCaller reduced total end-to-end processing time per sample by approximately 3.2-fold. In addition to faster performance, DupCaller provides greater flexibility by supporting Browser Extensible Data (BED) files, enabling users to restrict analyses to specific genomic regions. This feature is particularly advantageous for targeted duplex sequencing experiments, where mutation calling and rate estimation must be confined to predefined loci.

## DISCUSSION

This study introduces DupCaller, a probabilistic variant caller designed to improve the detection of somatic mutations from duplex sequencing data. Across synthetic benchmarks, mutagen-exposed cell lines, and normal tissue samples, DupCaller consistently identified more genuine somatic mutations than the NanoSeq analysis software, while maintaining high precision and accurately capturing known mutational signatures. Detecting more somatic mutations offers several important advantages: it reduces the likelihood of missing biologically relevant events— including potential driver mutations that may contribute to clonal expansion—and enables more accurate estimation of both mutation rates and mutational signatures’ activities across samples. In addition, DupCaller demonstrated faster computational performance and increased flexibility.

While DupCaller consistently identified more somatic mutations than the NanoSeq analysis software across all benchmarks, the magnitude of this difference varied between datasets. This variation arises from several factors. First, because DupCaller does not require a minimum number of reads per strand to call mutations, datasets with lower duplication rates and smaller read family sizes may benefit more from DupCaller compared to pipelines that enforce such thresholds. Second, each sample has a unique error profile, which can affect the sensitivity and specificity of mutation detection. In contrast, there are a small number of cases where DupCaller does not recover certain mutations reported by NanoSeq. These cases typically occur when mutations fall within regions with elevated error rates based on inferred sample-specific error profiles. A consistent example observed across benchmarks was single-base thymine insertions and deletions in long homopolymers. DupCaller’s probabilistic model assigns lower confidence to such contexts due to their high background error, while NanoSeq’s heuristic approach may still report them as true mutations. Since these are single-molecule events and cannot be directly validated, it remains uncertain whether DupCaller is being overly conservative or NanoSeq is reporting artifacts. However, given the known error-prone nature of these regions and the closer agreement between DupCaller’s indel burden and orthogonal expectations, it is more likely that these NanoSeq-specific calls represent artifacts.

Nonetheless, DupCaller has limitations. Its current implementation only supports data generated with barcode structures where equal-length barcodes are positioned at the start of each read—a design common to protocols like NanoSeq^15^, Duplex Sequencing^24^, BotSeq^16^, and EcoSeq^25^. As a result, DupCaller cannot yet process data from emerging technologies such as CODEC-seq^27^ and HiDEF-seq^30^, which use different barcode placements or sequencing platforms. However, the core algorithm is adaptable, and future updates may enable broader compatibility as these technologies gain traction. Additionally, reported benchmarks in this study focused on comparisons with the NanoSeq analysis software, as it represents the current state-of-the-art in duplex sequencing analysis. NanoSeq builds upon and systematically refines key aspects of earlier Duplex Sequencing and BotSeq protocols, including adapter design, ligation efficiency, consensus sequence generation, and filtering heuristics^15^. Its widespread adoption, demonstrated performance in recent high-profile studies, and the public availability of previously generated NanoSeq datasets make it the most relevant and rigorous comparator for evaluating DupCaller’s performance.

## ONLINE METHODS

### Overview of the DupCaller Algorithm

#### Sample preprocessing

The DupCaller pipeline takes paired-end FASTQ data as input, specifically generated with barcode structures in which equal-length barcodes are positioned at the start of each read. For each read pair, given the barcode structure of the reads, the barcodes and any universal bases are removed from the reads, and the barcode sequences of both reads are stored both in the read names and as tag “bc”. The sequences after barcode removal are then aligned to a reference genome of choice (*e.g.*, GRCh38 human genome) with BWA^39^ and the PCR duplicates and optical duplicates are identified with GATK MarkDuplicates^40^ with options *- umi -duplex*. For samples with an undiluted duplex sequencing data as matched control, the undiluted sequencing data are processed in the same way.

#### Error profile estimations

We considered two sources of errors to calculate confidence score of each base: single-strand mismatches and single-read discrepancies. Single-strand mismatches happen exclusively on one strand but not on the other strand. Single-read discrepancies are mismatches that are not supported by other reads of the same strand. Since a mismatch can be either a nucleotide mismatch or an insertion/deletion, four error matrices are generated per sample: single-strand SBS mismatches, single-strand indel mismatches, single-read SBS discrepancies, and single-read indel discrepancies. The estimation processes are as follow:

1. *Single-read discrepancy rate estimation:* For each single-strand read family with at least two reads on each strand, we first excluded any sites lacking the reference allele, as these are likely to represent true mutations. We also exclude sites that form non-reference consensus base on a single strand, as these can be derived from a single-strand mismatch. For the remaining bases, we count the occurrences of each of the four nucleotides given the trinucleotide context, weighted by their probability of sequenced correctly derived from the base quality score, resulting in a 64x4 matrix where the rows represent each of the 64 possible trinucleotide context and the columns represent each of the 4 nucleotides. For example, the entry [AAA] x [T] represents the number of sites with an AAA trinucleotide context and a reference base of A that were sequenced as T, indicating a potential error. In contrast, the entry at [AAA] × [A] counts sites with the same context where the reference base A was correctly retained. As we do not consider single-read discrepancies as stranded, each of the entry is added to its complementary entry (for example, [AAA] x [T] is added to [TTT] x [A]). For small insertions and deletions, instead of trinucleotide context, the error was categorized by the length of homopolymers preceding or proceeding the site and the length of the indel, generating a 40*21 matrix where the row represents the homopolymer length capped at 40 and the column represents the indel lengths are capped at −10 for deletions (loss of up to 10 base pairs) and +10 for insertions (gain of up to 10 base pairs), respectively.
2. *Single-strand mismatch rate estimation:* for each double strand family that has at least three reads on each strand, we excluded any bases altered in both strands and with discordant single-strand consensus, leaving only reference bases and bases that has concordant single strand consensus but discordant double strand consensus. The SBS and indel matrices are generated similarly to the single-read discrepancy estimation process. However, instead of aggregating complementary entries, sites on the bottom strand are normalized to their reverse complement. For example, a bottom-strand event recorded as [AAA] × [T] is treated as [TTT] × [A], whereas a top-strand [AAA] × [T] remains unchanged. For indels, strand normalization is not required, as neither the indel length nor the flanking homopolymer length is affected by strand orientation.

#### Genotype likelihood calculation

After estimation of context-specific error rates for each sample, we use the rates as parameters to genotype each consensus base using a probabilistic model (**Figure S7**). Since there are 16 possible nucleotide conversions, (*e.g.,* A>A, A>C, A>G, *etc.*) for both amplification and sequencing steps, complete posterior probability calculation requires calculating the posterior probability of all possible combinations of sequencing error and amplification error for all four bases and can be computationally intractable. We thus only consider a binary model in which only two alleles are considered for each locus within the same read family. If more than two alleles are present at the same locus, the locus is excluded for that read family. The proposed binary model (**Figure S6**) recapitulates the process of duplex sequencing library preparation. At a given locus within a duplex read family, we define the major base as B_1_ and the minor base as B_2_. If only the reference base is observed, B_2_ represents a composite base whose error rate is the average of the three possible non-reference nucleotides in the given sequence context. Conversely, if only a non-reference base is observed, B_2_ corresponds to the reference base. The observed random variables Q_1_ and Q_2_ are the summing base quality of bases on the top strand amplicons supporting B_1_ and B_2_ respectively, while N_1_ and N_2_ are the number of such bases. Similarly, Q_3_ and Q_4_ are the summing base quality of bases on the bottom strand amplicons supporting B_1_ and B_2_ respectively, while N_3_ and N_4_ are the number of such bases. The genotype likelihood of the duplex base being B_1_ is then computed as follows:

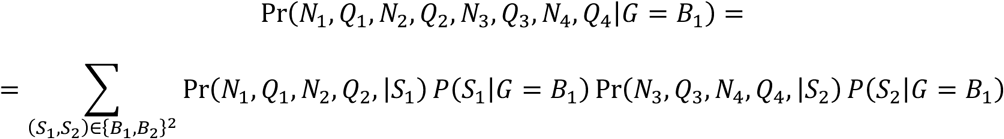

where

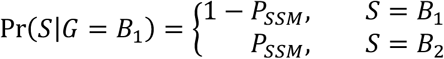

Here, P_SSM_ reflects the single strand mismatch rate of conversion from B_1_ to B_2_ at given trinucleotide context on top strand or bottom strand for S_1_ or S_2_, respectively.

Similarly, the genotype likelihood of duplex base being B_2_ is:

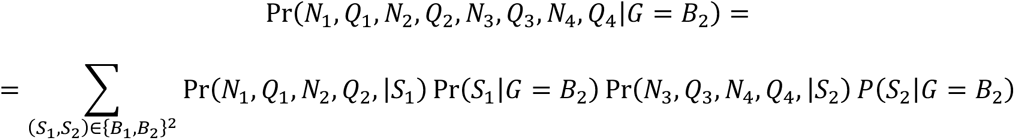

where

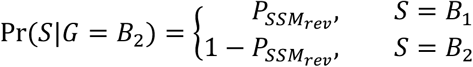

Here, P_SSM_rev_ is the single strand mismatch rate of conversion from B_2_ to B_1_ at the given trinucleotide context on top strand or bottom strand for S_1_ or S_2_, respectively.

For the top strand S_1_, the likelihood of S_1_ being B_1_ is:

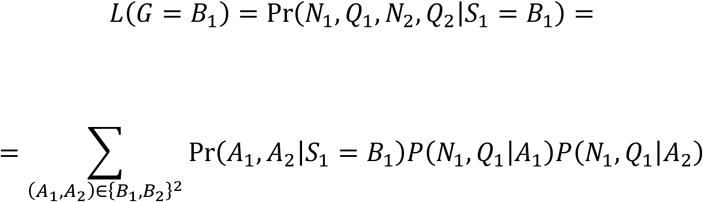

Modelling the single read discrepancies using a binomial distribution^41^, we get:

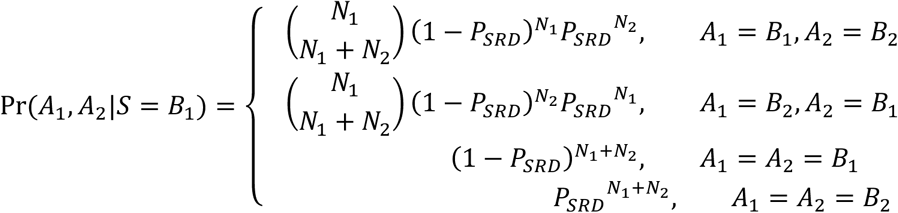

Here, P_SRD_ is the probability of single read discrepancy rate of conversion from B_1_ to B_2_ at the given trinucleotide context; and

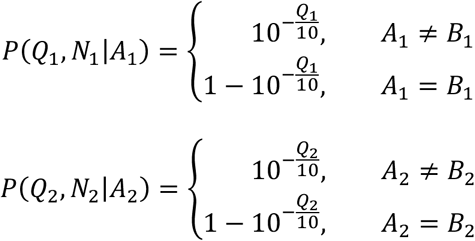

Similarly, we calculate likelihood of S_1_ being B_2_:

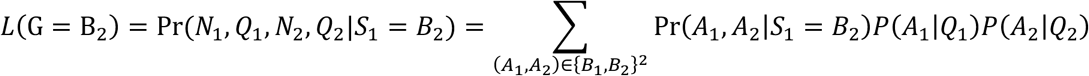

where

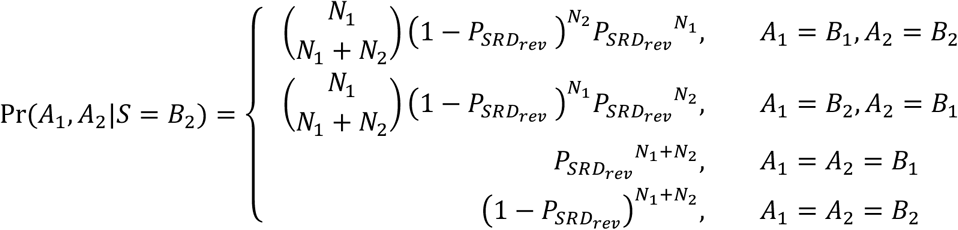

Here, P_SRD_rev_ is the probability of single read discrepancy rate of conversation from B_2_ to B_1_ at the given trinucleotide context. The likelihoods for the bottom strand S_2_ can be calculated analogously from Q_3_, Q_4_, N_3_, and N_4_.

We use the following confidence score (CS) to score the confidence to call the genotype as the major base B_1_:

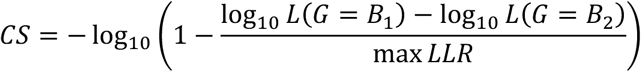

Where LLR is the log likelihood calculated previously and *maxLLR* is the maximum possible log likelihood ratio that happens when the single-read discrepancy rate is 0:

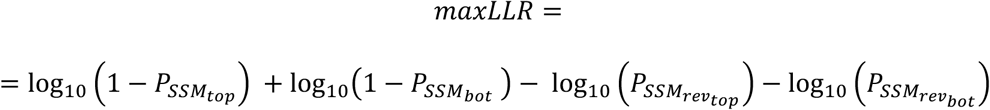

Mutations with a CS larger than a set threshold (default 0.50) were output as a candidate mutation. For indels, the CS calculation is similar with the exception that the top and bottom strand single-strand mismatch rate are equal.

#### Post-Likelihood Filtering Strategy

After mutation calling, we applied a set of read level and base level filters, similar to previous pipelines for processing duplex sequencing data^15,27^. Specifically, to ensure high-confidence variant detection, DupCaller applies both read-level and base-level filters. At the read level, it excludes reads with mapping quality below 50, as well as those marked as supplementary, secondary, optical duplicates, quality controlled failed, or with unsupported orientations. Reads with primary-secondary alignment score differences less than 50 are also filtered. Additionally, the number of mismatches in each read pair, calculated as the editing distance (NM) minus indel length, is calculated and any read pair with more than 5 mismatches are discarded. At the base level, positions with an allele frequency above 0.001 in a provided germline database are excluded; if no frequency is given, any site present in the database is omitted. Additional exclusions include bases masked by a noise mask, those lacking sufficient coverage in the matched normal sample, and mutations supported only by low-quality bases (i.e., base quality >18 in at least one read). For indels, an additional filter is applied using an indel-specific panel of normal samples constructed from whole-genome duplex sequencing datasets, which excludes sites where an indel appears in at least two samples with an allele frequency between 0.001 and 0.3. Reads failing read-level filters are excluded from mutation burden calculations, while sites failing base-level filters are omitted from burden estimation.

### Generation of synthetic datasets

Diluted and undiluted NanoSeq data of one cord blood samples from donor PD47269 (*donor A*) and 5 replicates of cord blood samples of donor PD48442 (*donor B*) were download from European Genome Archive with accession number EGAD00001006459. Germline mutations were called with GATK HaplotypeCaller^42^ and refined with VariantQualityScoreRecalibration. We then constructed a set of candidate mutations by taking germline mutations in *donor A* but not in *donor B*. For each replicate of *donor B*, mutations were randomly sampled from the candidate set and included in the ground truth if they met the following criteria: *(i)* the mutation was present in at least one duplex read family in the diluted *donor A* NanoSeq BAM file; *(ii)* no reads in the undiluted *donor B* BAM filed supported the mutation; and *(iii)* for indels only, if there does not exist other indels at the locus in undiluted donor B bam. Simultaneously, a random read family from *donor A* diluted NanoSeq that contains the ground truth mutation were added to the *donor B* diluted NanoSeq BAM to mimic a somatic mutation.

### Generation of genomic masks

Both DupCaller and NanoSeq analysis software utilize genomic masks of common germline mutations and noisy loci. For this study, we generated a noise mask, combining the original noise mask of NanoSeq with satellite regions near the centromeres of each chromosome, which we found to have a large portion of misaligned reads in NanoSeq data. For the germline mutation mask, we combined the NanoSeq single nucleotide polymorphism (SNP) mask with mutations with a population allele frequency larger than 0.1% based on gnomAD^43^ version 4.0.0. DupCaller also uses an indel panel of normal samples, as previously described. For the synthetic dataset, we generated a customized indel panel of normals by removing any sites overlapping with donor A germline mutations, to avoid excluding true positives from the ground truth set due to potential germline contamination in the original panel of normal.

### Benchmarking on synthetic dataset

DupCaller was run with default parameters except with the confidence score threshold set to 0.001 (-t 0.001) to output all candidate mutations, and synthetic dataset indel PoN. The precision and recall scores at each confidence score threshold were used to plot the precision-recall curve for DupCaller. NanoSeq analysis software (version 3.5.5) was ran with default parameters, except for minimum matched normal depth set to 10 (dsa -z 10) to match the corresponding parameter of DupCaller. Both pipelines use the *donor B* undiluted BAM as the matched control. The calling results for each pipeline was compared with the ground truth dataset to calculate the number of true positives (TP), false positives (FP), or false negatives (FN). False positive calls were further filtered by the noise and germline mask, and the precision, recall and F_1_ score was calculated.

For comparison of precision, recall and F_1_ score at default threshold, we corrected the number of FPs by the expected number of mutations in each PD48442 cord blood samples. Specifically, the number of FP is corrected as:

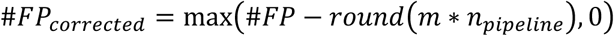

Where *m* is the expected mutation rate estimated from the whole genome sequencing from the cord blood colonies of the same donor (66 SBSs per cell, or SBS mutation rate of 1.2 x 10^-8^, and indel mutation rate of 6.5 x 10^-10^)^21^ and n_pipeline_ is the number of effective bases for each dataset using different pipelines. The corrected precision and F_1_ score were calculated based on the corrected number of FPs.

### Generation of mutagen exposed cell line datasets

HepG2 cell lines were purchased from ATCC (catalog #: HB-8065) and cultured per vendor’s recommendation. For the exposure experiment, the HepG2 cells were exposed to three different conditions: 40mM for 72 hours (AA_1), 70mM for 48 hours (AA_2), and 80mM for 48 hours (AA_3). Cells were cultured for one more passage after exposure and the harvested cell’s DNA was extracted with Qiagen DNeasy Blood and Tissue Kit (Catalog #: 69504). NanoSeq library preparation was done as previously described^15^, and sequencing was carried out using a NovaSeq 6000 sequencer.

### Benchmarking on cell line and human tissue NanoSeq data

NanoSeq data of 17 neurons and 7 cord blood samples were downloaded from European Genome Archive with accession number EGAD00001006459. DupCaller was run with default parameters, using the previously described genomic and noise masks. For indel calling, DupCaller also incorporated the full indel panel of unmatched normal samples. NanoSeq analysis software was run with default parameters, except for minimum control depth set to 10 (dsa -z 10) to match the corresponding parameter of DupCaller, and the same genomic and noise masks. The shared and caller-unique mutations were classified with BCFtools^44^ isec commands, and the mutation patterns were visualized with SigProfilerMatrixGenerator^31^. SBS mutation burdens were the corrected mutation reported by each pipeline. Indel burdens was calculated as the number of called indels divided by the number of effective bases for each pipeline.

### Benchmarking on sperm and blood saliva cohort

Sperm, blood, and saliva NanoSeq data was downloaded from dbGap with accession number phs003716.v1.p1. DupCaller was run on all samples same as previously have done, with the exception of extending the mutation calling regions to include chromosome Y. NanoSeq analysis software call sets and mutation rate estimations were gathered from the original publication^37^.

## SUPPLEMENTARY FIGURE LEGENDS

**Figure S1. Precision–recall performance of DupCaller and NanoSeq for single base substitutions on synthetic datasets.** Uncorrected precision–recall curves of DupCaller and precision–recall data points of the NanoSeq analysis software on synthetic single base substitution (SBS) datasets. The title of each panel indicates the number of ground truth mutations in that dataset; all datasets in the same row contain the same number of somatic SBSs. In each plot, the red line shows the DupCaller precision–recall curve, the red circle represents DupCaller’s default parameter setting, and the blue triangle marks the NanoSeq software result.

**Figure S2. Performance metrics of DupCaller and NanoSeq on synthetic benchmarking datasets.** Uncorrected performance metrics comparing DupCaller and the NanoSeq analysis software across 25 synthetic datasets, spanning five different ground truth mutation counts (five datasets per count). Metrics include uncorrected precision and uncorrected F_1_ score for ***(a)*** single base substitutions (SBS) and ***(b)*** insertions/deletions (indels). Bars are color-coded by the number of spiked-in ground truth mutations. For each mutation count, the bar height represents the average value across replicates, and the error bar indicates the standard deviation.

**Figure S3. Precision–recall performance of DupCaller and NanoSeq for small insertions and deletions on synthetic datasets.** Uncorrected precision–recall curves of DupCaller and precision–recall data points of the NanoSeq analysis software on synthetic insertion and deletion (indel) datasets. The format and layout mirror those of *Figure S1*, with each panel corresponding to a specific number of ground truth indels. The red line represents the DupCaller precision–recall curve, the red circle indicates DupCaller’s default parameter setting, and the blue triangle denotes the NanoSeq result.

**Figure S4. Effect of duplication rate on relative mutation detection by DupCaller.** This figure shows the ratio of single base substitutions (SBSs; y-axis) identified by DupCaller compared to those identified by the NanoSeq analysis software, plotted against the duplication rate for each sample (x-axis). Samples include those from the aristolochic acid–exposed HepG2 cell line, neurons, and cord blood. The data illustrate how lower duplication rates are associated with greater gains in mutation detection by DupCaller, highlighting its ability to recover more mutations under suboptimal sequencing conditions.

**Figure S5. Comparison of SBS and indel mutational spectra in cord blood samples.** Mutational spectrum analysis of seven cord blood samples comparing variant calls from DupCaller and the NanoSeq analysis software. ***(a)*** SBS-96 profiles for mutations detected by both tools, as well as those uniquely identified by DupCaller or NanoSeq. ***(b)*** Corresponding indel spectra shown using the ID-83 classification for the same shared and tool-specific call sets.

**Figure S6. Comparison of computation time between DupCaller and NanoSeq analysis pipelines**. Bar plots show the average computation time (y-axis) for equivalent processing steps in DupCaller and NanoSeq analysis software pipelines (x-axis), using 17 neuron samples. Each bar represents the average time per step, with error bars indicating standard deviation across the eight samples. The “tag” step is specific to the NanoSeq pipeline and is not used by DupCaller.

**Figure S7. Strand-specific probabilistic model for genotype inference from duplex sequencing data.** The model assumes that each locus within a read family contains only two candidate nucleotides: a major allele (*B*_1_) and a minor allele (*B*_2_). The true underlying genotype *G* ∈ {*B*_1_, *B*_2_} is associated with a top strand nucleotide (S1) and a bottom strand nucleotide (S2), each modeled as a categorical distribution conditioned on *G*. For each strand, *A*_1_ represents the amplicons of the top strand sequenced into *N*_1_ ∈ ℤ reads supporting base *B*_1_, with a consensus base quality *Q*_1_ ∈ ℤ. Similarly, *A*_2_ represents the amplicons supporting base *B*_2_, sequenced into *N*_2_ ∈ ℤ reads with quality *Q*_2_ ∈ ℤ. The model is used to compute the likelihood of observed data given candidate genotypes. Full details of the probability calculations are provided in the *Methods* section.

## DATA AVAILABILITY

The aristolochic acid exposure experimental datasets have been deposited in the Sequence Read Archive (SRA) under project accession number PRJNA1262723. The cord blood and neuron samples were download from European Genome Archive with accession number EGAD00001006459. The sperm, blood, and saliva cohort can be downloaded from dbGap with accession number of phs003716.v1.p1. The synthetic benchmarking dataset is based on samples from EGAD00001006459 and can be reproduced using the code provided with this manuscript.

## CODE AVAILABILITY

DupCaller is an open-source tool released under the permissive BSD-2 software license. The complete source code, along with documentation and example usage, is freely available on GitHub at https://github.com/AlexandrovLab/DupCaller.

The code used to generate the synthetic benchmark dataset is available on GitHub at https://github.com/AlexandrovLab/DupCaller_manuscript and can be utilized to generate a synthetic benchmark dataset when applied to the duplex sequencing data from European Genome Archive with accession number EGAD00001006459.

## ACKNOWLEDGMENTS

L.B.A. and Y.C. would like to thank Federico Abascal and Iñigo Martincorena for their valuable discussions and insightful advice during the development of DupCaller. This work was supported by the US National Institute of Health grants R01ES032547, R01ES036931, R01CA269919, R01CA296974, P01CA281819, and U01CA290479 to L.B.A. as well as by L,B.A.’s Packard Fellowship for Science and Engineering and the UC San Diego Sanford Stem Cell Institute. The computational analyses reported in this manuscript have utilized the Triton Shared Computing Cluster at the San Diego Supercomputer Center of UC San Diego. The funders had no roles in study design, data collection and analysis, decision to publish, or preparation of the manuscript.

## AUTHOR CONTRIBUTIONS

Y.C. and L.B.A. conceptualized the study and designed the DupCaller algorithm with assistance from S.P.N. Y.C. implemented the DupCaller algorithm and performed benchmarking with assistance and advice from S.P.N., S.A., L.C., and M.P. S.P.N performed aristolochic acid exposure experiments with assistance from A.K. and I.S. Y.C. and L.B.A. wrote the manuscript with input from all co-authors. All authors read and approved the final manuscript.

