## Supplementary figures and images for "Improved Mutation Detection in Duplex Sequencing Data with Sample-Specific Error Profiles"

### Supplementary Figures S1-S7

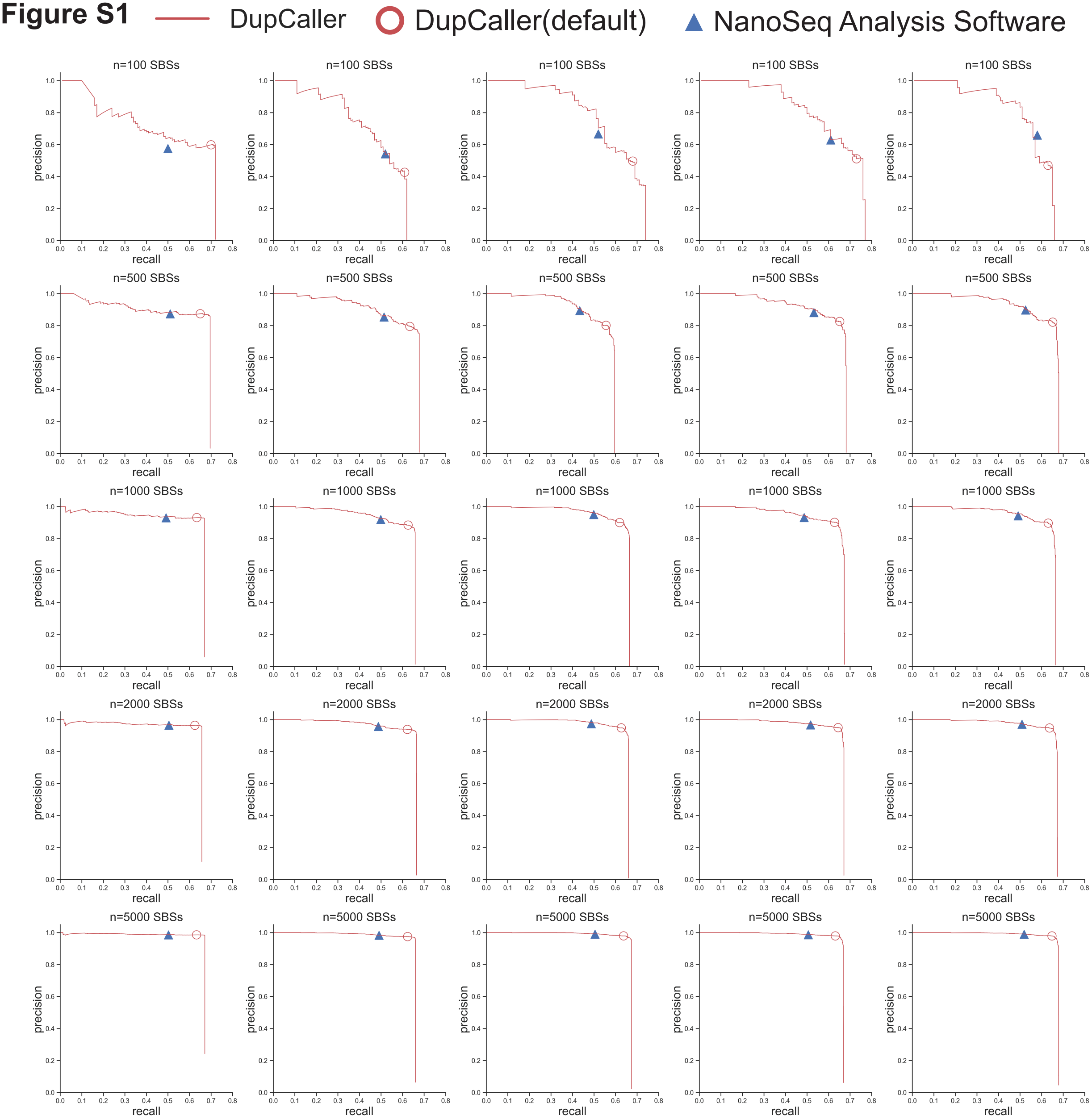

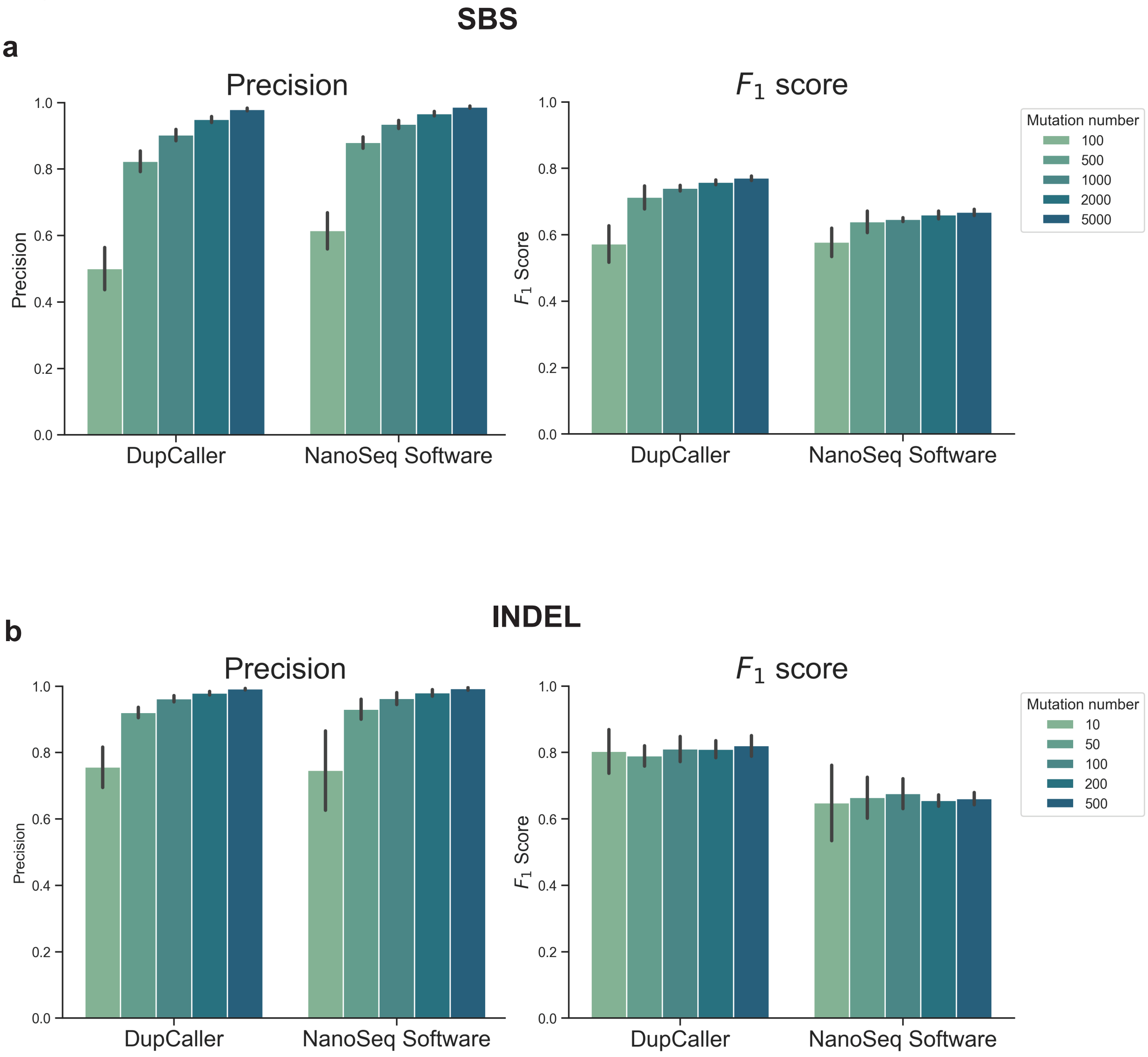

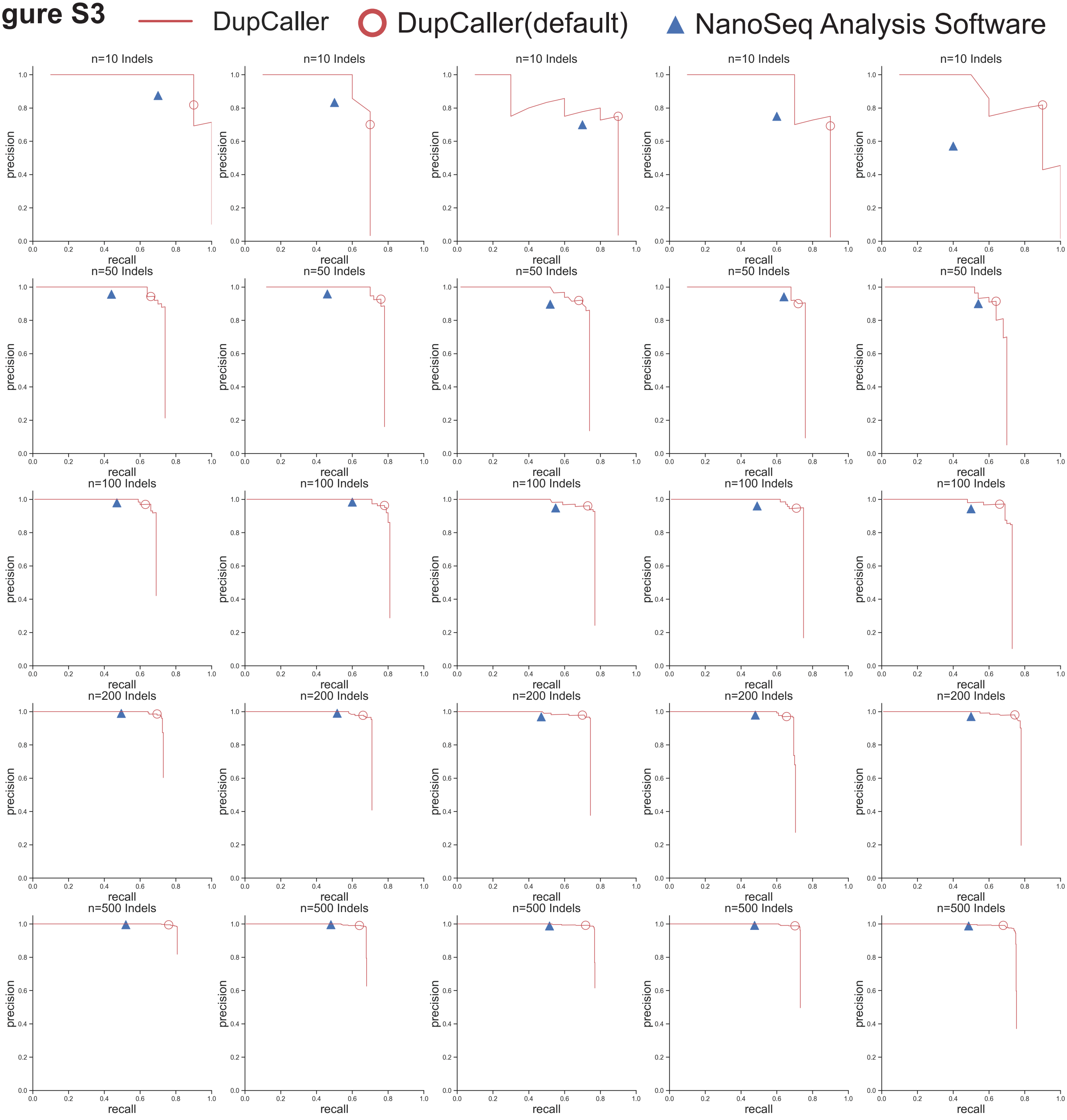

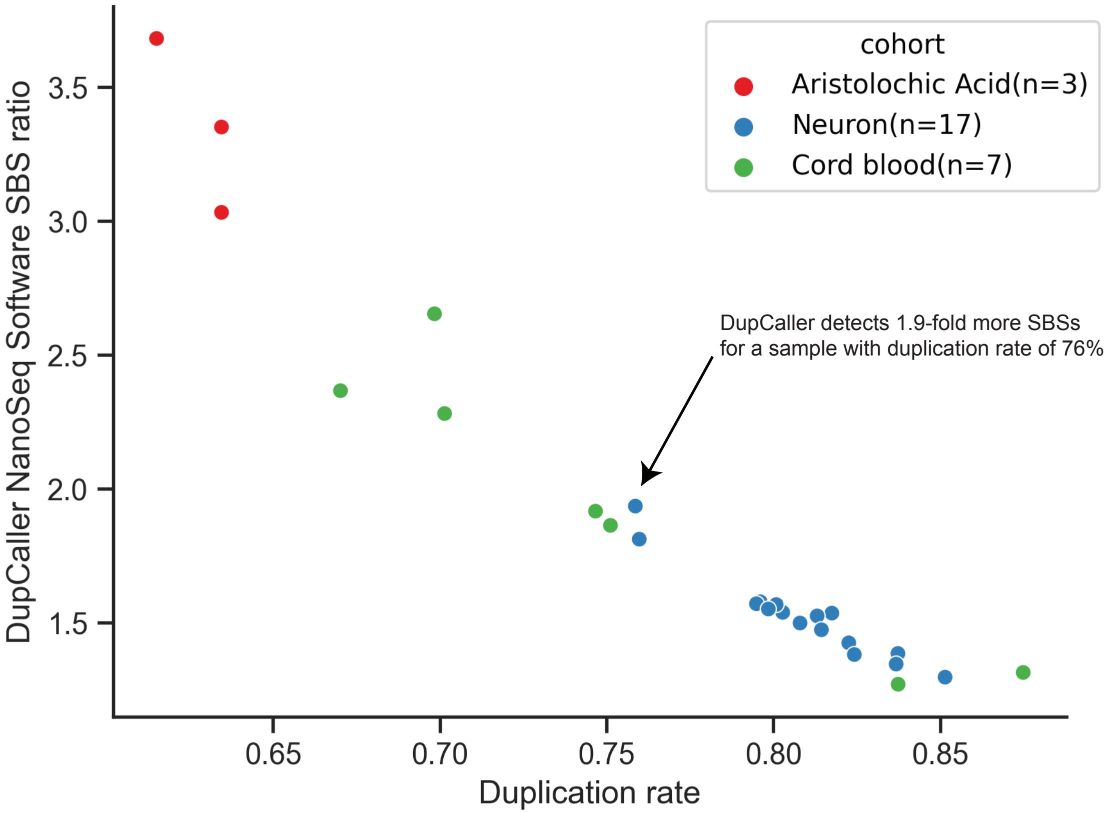

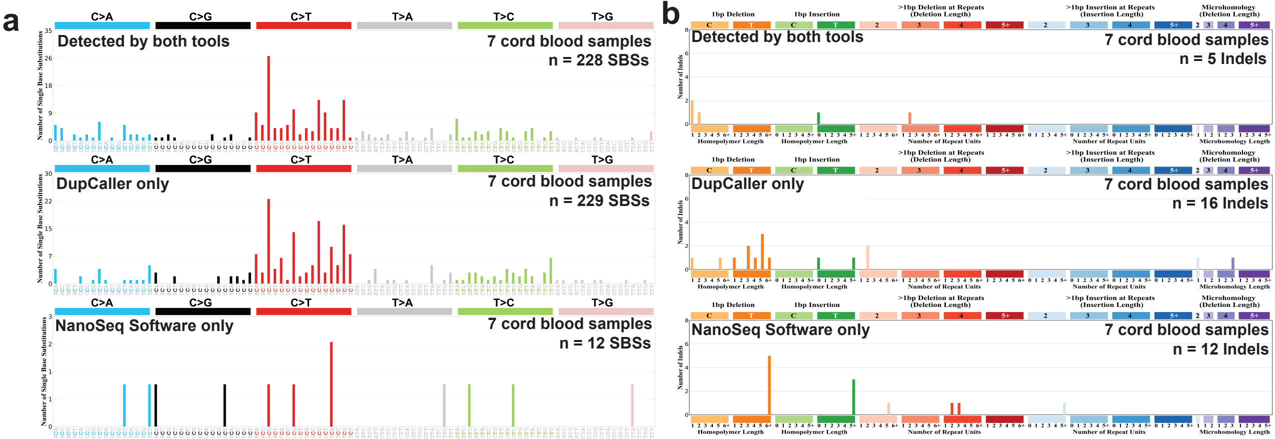

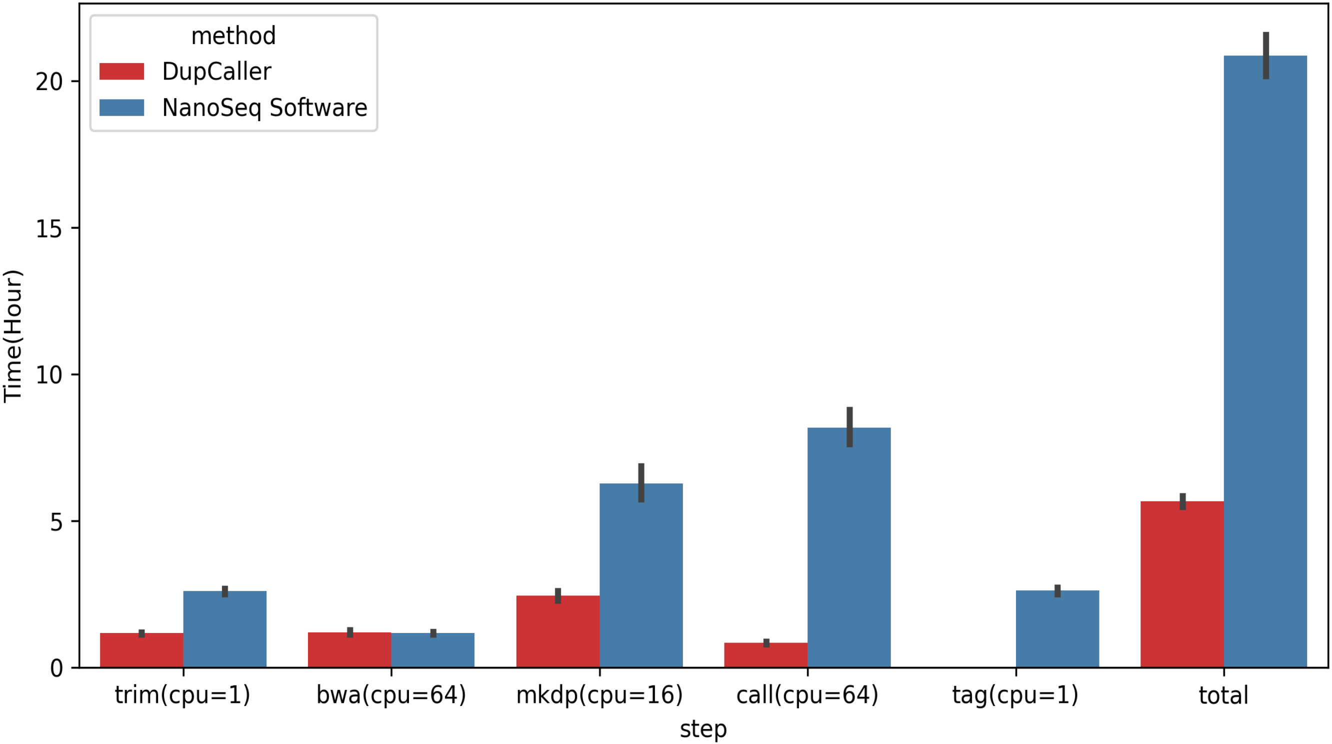

Figure S7

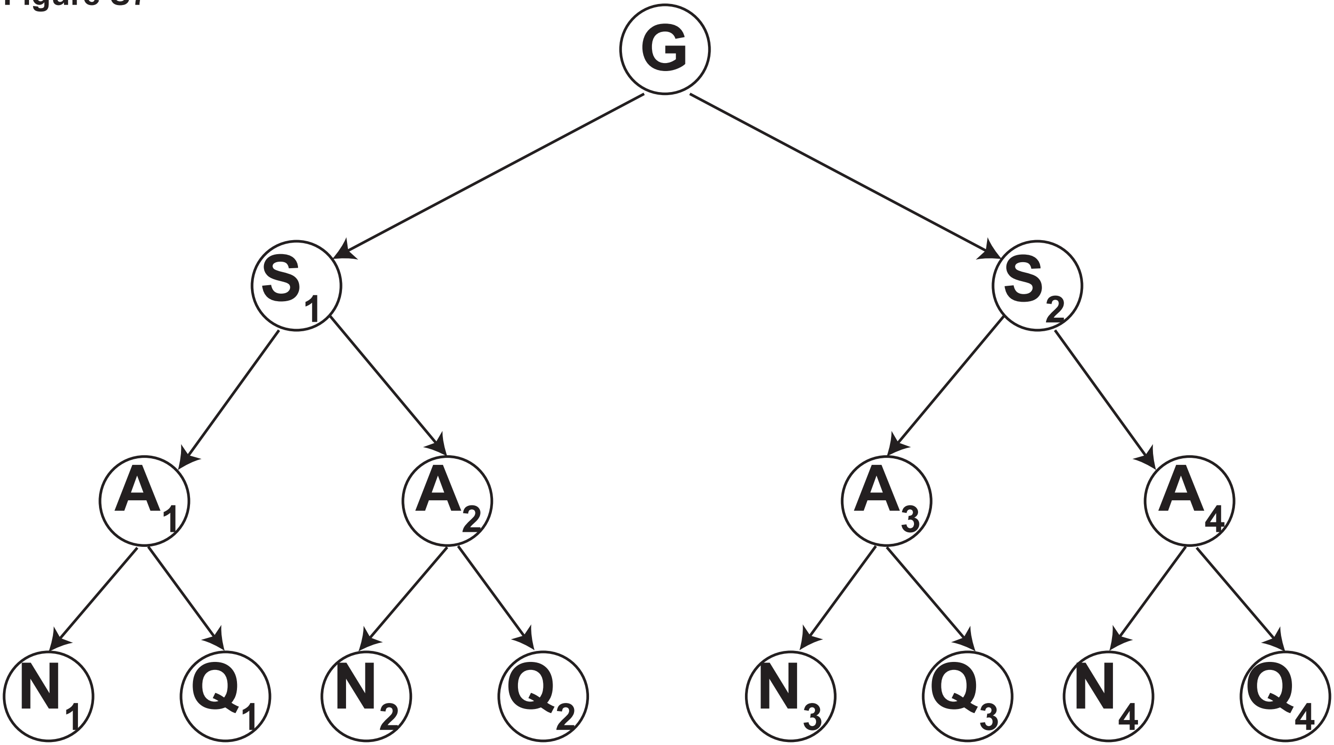
